# SERS nanowire chip and machine learning enabled instant identification and classification of clinically relevant wild-type and antibiotic resistant bacteria at species and strain level

**DOI:** 10.1101/2023.01.12.523744

**Authors:** Sathi Das, Kanchan Saxena, Jean-Claude Tinguely, Arijit Pal, Nima L. Wickramasinghe, Abdolrahman Khezri, Vishesh Dubey, Azeem Ahmed, Perumal Vivekanandan, Rafi Ahmad, Dushan N. Wadduwage, Balpreet Singh Ahluwalia, Dalip Singh Mehta

## Abstract

The world health organization considers antimicrobial resistance (AMR) to be a critical global public health problem. Conventional culture-based methods that are used to detect and identify bacterial infection are slow. Thus, there is a growing need for the development of robust, cost-effective, and fast diagnostic solutions for the identification of pathogens. Surface-enhanced Raman spectroscopy (SERS) can be used to identify target analytes with sensitivity down to the single-molecule level. Here, we developed a SERS chip by optimizing the entire fabrication pipeline of the metal-assisted chemical etching (MACE) method. The MACE approach offers a large-scale, densely packed silver (Ag) nanostructure on top of silicon nanowires (Si-NWs) with a large aspect ratio that significantly enhances the Raman signal due to localised surface plasmonic enhancement. The optimised SERS chips exhibited sensitivity down to 10^-12^ M concentration of R6G molecule and detected reproducible Raman spectra of bacteria down to a concentration of 100 colony forming units (CFU)/ml, which is a thousand times lower than the clinical threshold of bacterial infections like UTI (10^5^ CFU/ml). A Siamese neural network model was used to classify SERS Raman spectra from bacteria specimens. The trained model identified 12 different bacterial species, including those which are causative agents for tuberculosis and urinary tract infection (UTI). Next, the SERS chips and another Siamese neural network model were used to differentiate antibiotic-resistant strains from susceptible strains of *E. coli*. The enhancement offered by SERS chip enabled acquisitions of Raman spectra of bacteria directly in the synthetic urine by spiking the sample with only 10^3^ CFU/ml *E. coli*. Thus, the present study lays the ground for the identification and quantification of bacteria on SERS chips, thereby offering a potential future use for rapid, reproducible, label-free, and low limit detection of clinical pathogens.

## Introduction

In recent years, the massive increase of infections caused by microorganisms has sought immediate attention from the public health care sector.^1,2^ Microorganisms can cause severe infections and cause significant morbidity and motality.^2^ According to WHO, more than 1.2 million people died from severe bacterial infections in the respiratory, gastrointestinal, and central nervous systems.^3^ A healthy individual can acquire pathogens either from other humans, the environment (contaminated food, water, and air) or pets, farm animals (zoonotic transmission).^4–8^ Thus, there is an ardent need to develop a fast, accurate detection method to diagnose bacterial infections. The traditional detection techniques, including polymerase chain reaction (PCR), flow cytometry, and enzyme-linked immunosorbent assay (ELISA), are time-consuming, laborious, and require specific laboratory infrastructure and trained personnel. Thus, the culture-based detection technique precludes its use for rapid, ultralow detection of pathogens.^8^ Furthermore, regular exposure to antibiotics often drives the evolution of resistance via horizontal or vertical transfer of resistant genes, thereby enabling bacteria to escape antibiotic assaults. Consequently, the treatment of multidrug-resistant pathogens is becoming a challenge for clinicians. Thus, it is of utmost importance to determine the phenotypic characteristics of bacterial strains for rapid clinical diagnosis and initiation of a proper treatment regime in time.

Raman spectroscopy is a popular molecular fingerprinting technology that enables non-destructive, sensitive, and quantitative detection of molecules in terms of their vibrational and rotational energy levels.^9^ However, Raman scattering is an intrinsically weak phenomenon. Consequently, it requires a very dense sample or a high laser input to achieve a valuable peak intensity in the spectrum. Thus, this technique with a low laser power is not compatible with ultrasensitive detection.^10^ Surface-enhanced Raman spectroscopy (SERS), enhances the weak traditional Raman signal using the nanostructure of silver (Ag) or gold (Au).^11–13^ The Ag and Au nanoparticles exhibit plasmonic resonance in the visible range of light, enabling the trapping of light at the nanoscale and further enhancing the local electric field. As a result, when an analyte is placed over the enhanced local electric field, the scattered Raman signal is amplified. The SERS as a detection technique is becoming popular in biomedical, and clinical fields for routine analysis of diseases.^14^ For example, Dina et al. have reported the detection and identification of pathogens using Ag nanoparticles-based SERS substrates.^15^ Boardman et al. have reported an Au nanoparticle coated SiO2 SERS substrate for detecting pathogens in human blood.^16^

The nanostructures of anisotropic shapes and sizes offer a greater amount of local electric-field enhancement compared to the nanoparticles with a spherical shape.^13^ Especially, the three-dimensional nanostructures with sharp tips and closely spaced nanostructures exhibit extremely high induced electric field at the sharp edges (rod lightening effect), and the enhancement due to plasmonic coupling resonance hybrid modes.^17^. To generate periodic nano-patterns lithography methods like electron beam lithography, nanoimprint lithography techniques are usually employed for fabricating SERS substrates. However, lithography techniques are time-consuming, costly, and electron beam is unsuitable to produce large chip-size and for mass production.^18^ Thus, there is continued research interest in the development of alternative fabrication process for affordable and scalable SERS-chip. An alternative fabrication route is metal-assisted chemical etching (MACE) process, which is a facile, scalable, and reproducible technique for fabricating high aspect ratio Si nanowires.^19^ MACE technique allows fabrication of highly structured Ag-coated Si nanowire assays over a large surface area. The MACE technique is also advantageous since it does not require any expensive vacuum equipment like other physical or chemical vapour deposition methods. Several publications report the scalability, reproducibility, and sensitivity of SERS substrates fabricated by the MACE technique for the ultrasensitive detection of explosive molecules, toxic chemicals, etc.^20–22^ However, only a few research report the detection of biomolecules and microorganisms using an Ag-coated Si nanowire SERS substrate^23,24^. Karadan et al. have reported a lithography-modified Ag-coated Si nanopillar-based architecture as a SERS substrate for detecting *E. coli* bacteria and malaria-infected red blood cells.^23^ Daoudi et al. have detected the spike protein of the SARS CoV-2 virus using an Ag nanoparticle/Si NW SERS substrate.^24^ However, the systematic detection of pathogens at the species and the strain levels using an Ag-coated Si NW SERS substrate has not been systematically investigated previously.

In this work, we developed and optimized the SERS chip using MACE technology and demonstrated its utility for label free detection of Raman spectra from twelve different strains of non-pathogenic and pathogenic bacteria and of two antibiotic-resistant strains of *E. coli*. To detect and classify Raman spectra from SERS chips, we then trained a Siamese neural network model. Along with the model, the SERS chips detected Raman signatures of bacteria even at a low concentration of 100 colony forming units (CFU)/ml. The Raman spectra from different bacteria were classified at species and strain levels using the trained model. For species level classification, we have selected a diverse set of bacteria having different cellular morphology, gram-staining property, and spore-forming capability. Further, for strain level classification, SERS spectra were generated for three different strains of *E. coli* including antimicrobial-resistant (AMR) strains that were validated using whole genome sequencing.^25–27^ The study emphasizes the utility of the fabricated SERS chip and the neural network model in performing a label-free, real-time, and molecular-specific detection of wide range of bacteria, enabling it to be advantageous over existing culture-based diagnostics approaches.

## Methods and Materials

### Fabrication and optimization of SERS chip

We have used a facile two-step MACE fabrication method to produce silver (Ag) coated Si nanowire (SiNW) SERS chips. The MACE method has three essential requirements: 1. Hydrofluoric acid (HF) as a complexing agent; 2. Ag nanoparticles for catalysis (reduced from silver nitrate, AgNO_3_); and 3. an oxidant with a higher reduction potential than silicon (hydrogen peroxide, H_2_O_2_). The fabrication principle, schematized in Figure 1, starts with the wet-chemical deposition of silver on the silicon surface. In a next step, H_2_O_2_ is reduced with the metal as a catalyst. The created protons inject electronic holes into the Si close to the silver particles, where the electrolyte locally oxidizes the substrate to SiO2. SiO2 is then etched by HF, where the sinking metal particles during the ongoing etching process generate pores with high aspect ratio leaving behind nanopillar-like structures, as shown in Figure 2 (a). The subsequent removal and re-coating of Ag nanoparticles on the SiNW is required for exploring the optimized plasmonic property of such metallic nanostructures in SERS applications. The last step with silver redeposition (Figure 2 (b)) was optimised for maximum SERS performance considering a 1 micromolar (10^-6^ M) concentration of Rhodamine 6G. The detailed experimental protocol, substrate morphology study, substrate optimization and numerical simulations are described in Supplementary sections 1-4, with more information on the MACE principle to be found in literature.^20^

**Figure 1.**
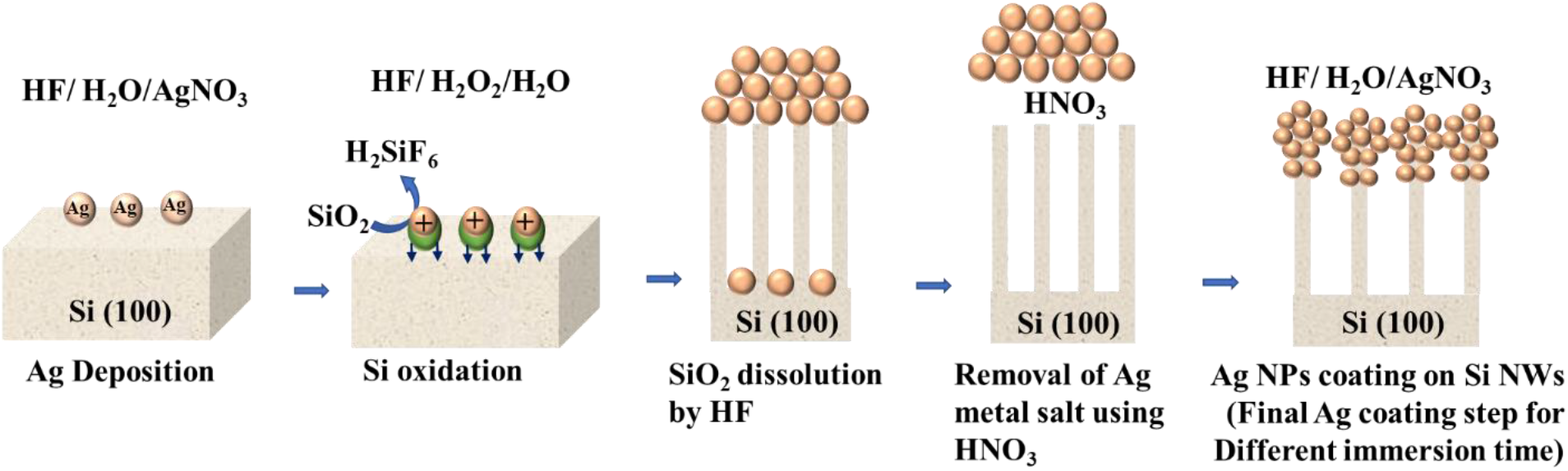
Schematic diagram of the fabrication process of the utilized SERS substrate by means of the wet-chemical MACE technique. First, silver nanoparticles are deposited on the silicon surface. In a following step, H_2_O_2_ oxidation is catalyzed by the metal surface, which locally oxidizes silicon allowing for its removal with HF. The ongoing etching process leads the silver particles to sink into the silicon crystal, creating nanopillars with high aspect ratio. After a first removal of the silver particles, another deposition step provides an ideal distribution around the nanopillars producing inexpensive and large area SERS substrates.

**Figure 2.**
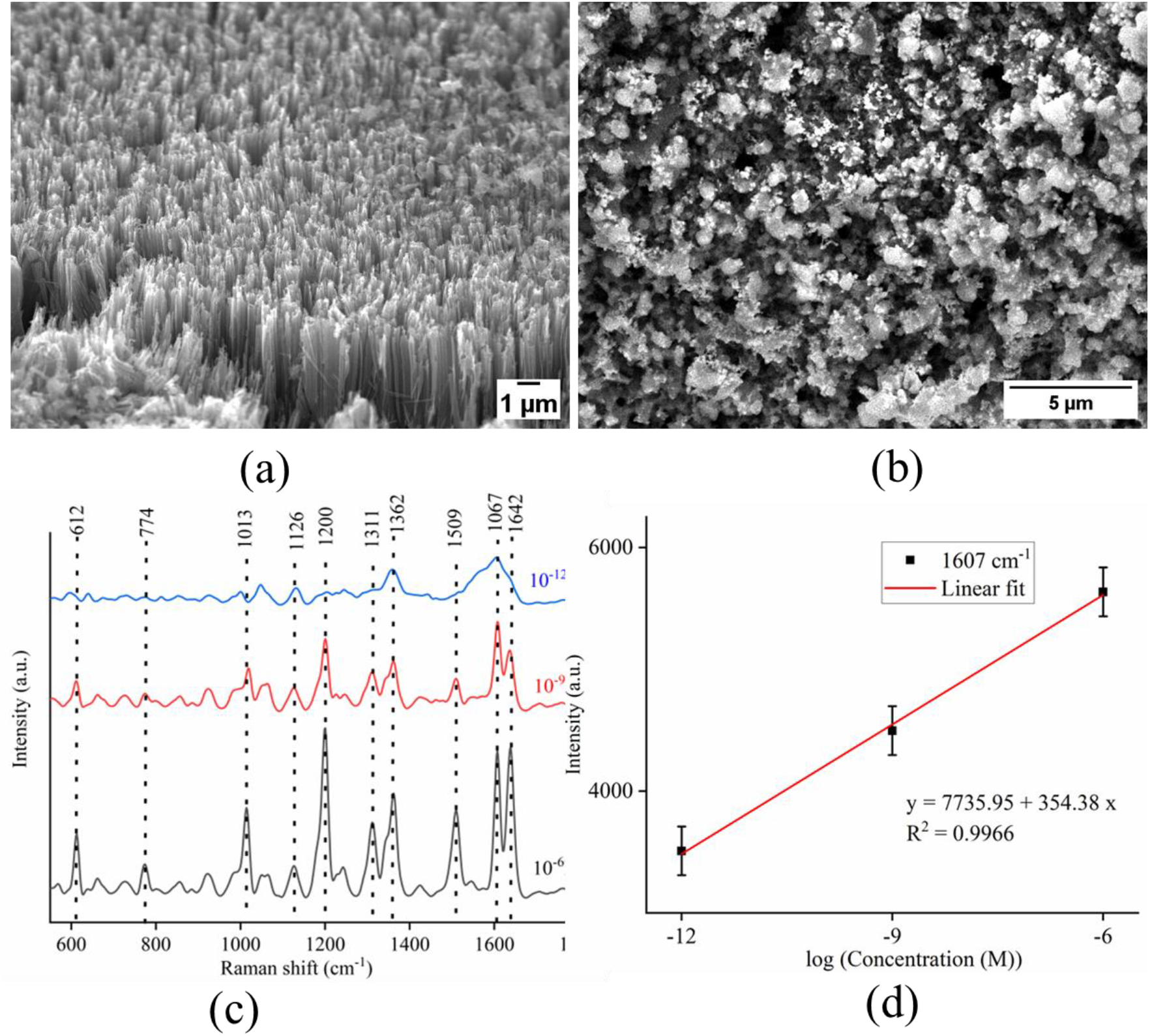
SEM images of (a) SiNW, (b) Ag coated Si NWs,Morphology, where (a) shows tilted view, and (b) shows top view of the nanostructure,(c) Chip sensitivity by considering the SERS spectra of R6G for micro -pico molar concentrations, (d) Linear dependence of signal intensity vs. concentration for a selected Raman peak.

Figure 1 shows the schematic diagram of the fabrication process of the SERS chip. The details of nanostructure fabrication, morphology of nanostructures and optimisation of Ag coated Si NW SERS chip is provided in the supplementary information at Supplementary Section 1-3 and supplementary figures, S.F. 1-4 respectively. Figure 2 (a,b) presents scanning electron microscopy (SEM) images of the SiNW surface, before (a) and after (b) metallic Ag deposition, respectively.

### Model architecture for machine learning

#### Classification strategy

We developed the workflow to specifically address two aspects that are suitable for complex real-life situation: can a classification model generate valid classification data from a limited number of samples; and secondly is it possible to reliably classify distinct bacteria species regardless of concentration.

To fix the limitation occurring due to the limited number of samples, a Siamese model was utilized for classification. Siamese models have proven effective with small sample sizes.^28^ Two inputs are required for classification using a Siamese model. The model outputs whether these two inputs are similar or not. A base set was created for this purpose by taking one spectrum from each class of bacteria. Each input is compared to the SERS spectra of seven class of bacteria (two tuberculosis causing bacteria and 5 UTI-causing pathogens) in the base set, and the signal with the highest similarity score is selected as the predicted class. The details of model architecture is discussed in the Supplementary Section 5 and shown in supplementary figure S.F. 5.

### Results and discussions

Post-optimization, the reproducibility of the spectrum from SERS substrate was investigated considering different spots on a single SERS substrate and substrates from multiple fabrication batches, as shown in Supplementary Figure S.F. 2 (b, c), respectively. The Raman peaks are highly reproducible, and the standard deviation of peak intensity is found to be less than 10%. Thus, the performance of the optimised SERS substrate is reproducible in different batches and at different locations within a single chip. The SERS activity was further checked by considering the detection limit, taking R6G as the probe molecule. Figure 2 (c) shows that the optimised SERS chip exhibits enhanced SERS spectra up to picomolar (10^-12^ M) concentration of R6G. Figure 2 (d) gives the Intensity vs concentration plot curve taking three different concentrations (10^-6^ M to 10^-12^ M) of R6G considering a particular peak at 1607 cm^-1^ for 10 measurements over random spots. The calibration plot fits linearly and matches well with the experimental points, as shown in Figure 2 (d). The plot shows the linear variation of a particular Raman peak with concentration. Table 1 depicts the vibrational bands associated with R6G molecule.

**Table 1.** Assigned SERS spectrum peaks of R6G ^29,30^.

| Raman peaks (cm<sup>-1</sup>) | Assigned vibrational peaks |
| --- | --- |
| 612 | In plane bending in C-C-C ring |
| 774 | Out of plane bending of C-H |
| 1013 | Hybrid modes of xanthene and phenyl rings |
| 1126 | In plane bending C-H |
| 1200 | In plane bending of C-H in xanthene ring |
| 1311 | C-O-C in plane bending |
| 1362, 1509, 1607, 1642 | Aromatic C-C stretching |

### SERS chips for acquisitions of Raman spectra from different species of bacteria

Post-optimisation, we acquired SERS spectra of individual bacteria solution using Ag coated Si NWSERS chips and created a database of vibrational energy levels incorporating molecular fingerprint information. The SERS spectra were collected from 12 groups of disease-causing microbes: *Mycobacterium tuberculosis* H37Ra (ATCC 25177), *Mycobacterium smegmatis* MC^2^155 (ATCC 700084), *Escherichia coli* K12 (ATCC 700926), *Klebsiella pneumoniae* 33495 (ATCC 33495), *Enterobacter cloacae* 10005 (ATCC 13047), *Pseudomonas aeruginosa* Migula (ATCC 27853), *Staphylococcus aureus* 7447 (ATCC 6538P), *Paenibacillus borealis* SB1 (MTCC 8085), *Bacillus clausii* 1779 (MTCC 11713), *Bacillus subtilis* 3610 (MTCC 121), *Xanthobacter autotrophicus* 10809 (MTCC 132), and *Pseudomonas citronellolis* 50332 (MTCC 1191). The bacterial sample preparation details are provided in the supplementary section 6. The assigned vibrational Raman peaks of bacteria are listed in Table 2. Figure 3 displays the SERS spectra of various bacteria having 10^5^ CFU/ml concentration, which is the clinically accepted concertation for a bacterial infection. Below we discuss each sub-groups shown in Fig. 3 separately.

**Table 2.** Assigned SERS spectrum peaks acquired for bacteria^31–33^.

| Raman bands | Assigned vibrational peaks | Biochemical Group |
| --- | --- | --- |
| 644-657 | C-S stretching, C-C twisting Guanine (C-S) | Proteins and nucleic acids |
| 674 | $\nu$(C-S) Guanine | Nucleic acids |
| 724-738 | $\rho$(CH<sub>2</sub>) Adenine, trans conformation of (C-S) or tryptophan | Nucleic acids |
| 850-855 | $\nu$(CC) | Proteins |
| 1001-1005 | $\nu$ (CC), aromatic ring of Phenylalanine | Protein |
| 1057-1061 |  | Proteins, Nucleic acids |
| 1079 | Phenylalanine | Protein |
| 1124-1126 | Carbohydrates, C-N and C-C stretching | Polysaccharide |
| 1180-1183 | C-C and C-H stretching | Lipids |
| 1242-1248 | Cytosine, adenine (DNA), amide III | Nucleic acids and Proteins |
| 1282 | CH deformation (proteins) | Proteins |
| 1321-1333 | $\delta$ (CH)Adenine | Nucleic acids |
| 1369 | (C-H<sub>2</sub> deformation or tryptophan) | Nucleic acids |
| 1471-1472 | $\delta$ (C-H<sub>2</sub>) protein | Protein |
| 1573 | $\nu$(C-N), $\delta$(N-H) of amide II | Protein |
| 1646 | amide I α- helix | Protein |

**Figure 3.**
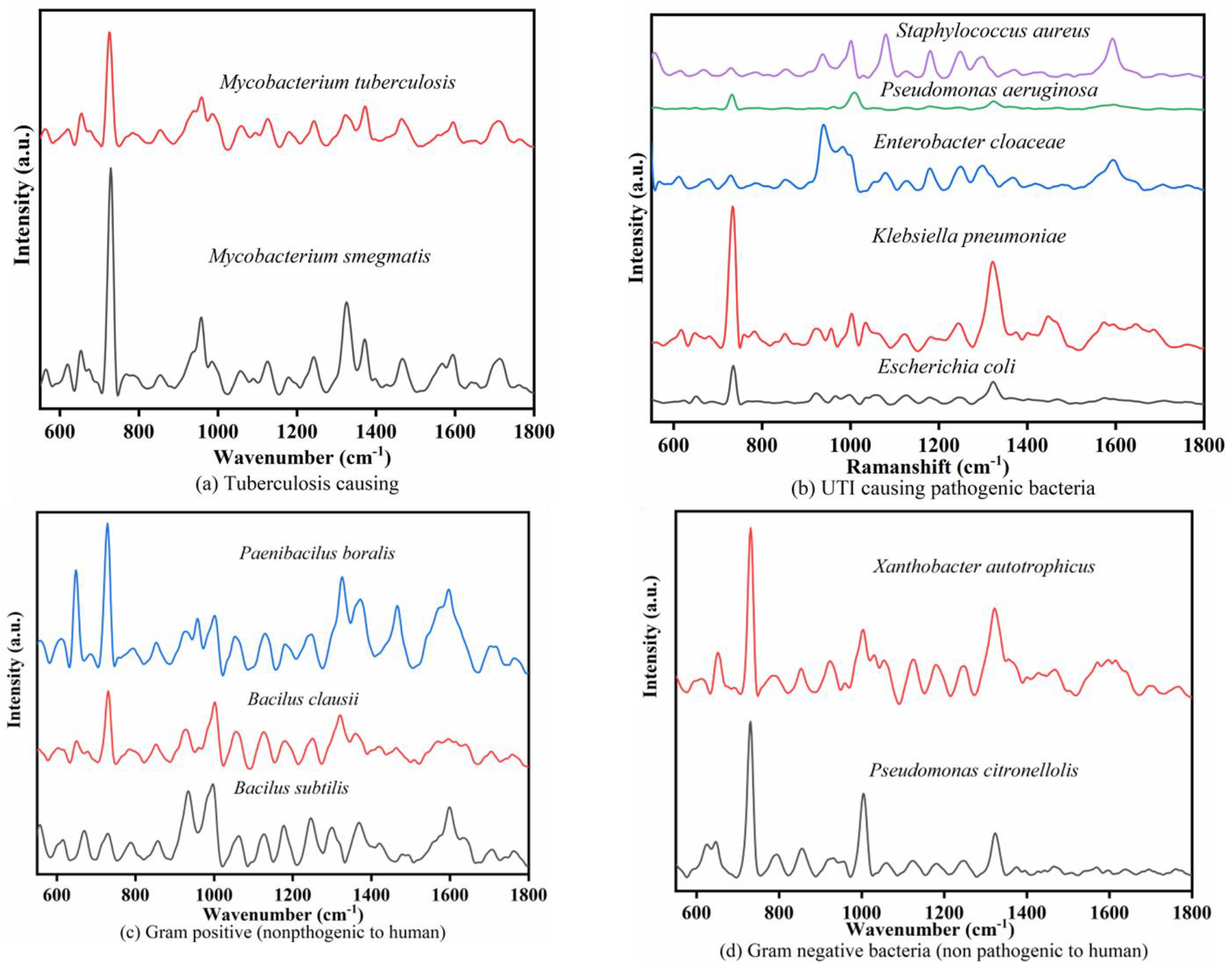
SERS spectra SERS spectra of 12 different classes of bacteria having concentrations of 10^5^ CFU/ml. The groups are formed based on the causing diseases. (a) Tuberculosis, (b) Urine track infection (UTI), two gram straining properties based on plant pathogenic bacteria: (c) gram positive, (d) gram negative.

Figure 3(a) shows that the SERS spectral features of two tuberculosis causing bacteria, Mycobacterial species-*Mycobacterium smegmatis* and *Mycobacterium tuberculosis H37Ra* are similar. Perumal *et al* have shown that the SERS characterisation of Mycolic acid (MA) as a TB biomarker because of their high abundance (up to 50% of bacterial dry weight is composed of MA) and its stable and inert nature.^31^ MAs are long-chain fatty acids that serve as a diagnostic biomarker for mycobacterial infection. The SERS spectra exhibits distinct features of intact MAs showing peaks at 654, 724, 852, 985, 1057, 1124, 1242, 1321,1371, 1466, 1592, and 1706 cm^-1^. The assigned vibrational modes are described in Table 2. However, *MTB H37Ra* displays some additional Raman peaks as compared to that from *Mycobacterium smegmatis*, indicating the structural complexity of the human pathogen.

Figure 3(b) shows the SERS spectral features of 5 UTI-causing pathogens *Escherichia coli* K12 (ATCC 700926), *Klebsiella pneumoniae* 33495 (ATCC 33495), *Enterobacter cloacae* 10005 (ATCC 13047), *Pseudomonas aeruginosa* Migula (ATCC 27853), and *Staphylococcus aureus* 7447 (ATCC 6538P). The Raman spectra from these bacteria exhibit a few common and many distinct spectra features as shown in Figure 4 (b). The peak at 735 cm^-1^, 1323 cm^-1^, and 1124 cm^-1^ are the strongest and common in all the species. These highly intense peaks prevailing the binding of the Ag nanoparticles with the bacteria cell wall are attributed to in-plane ring breathing mechanism of adenine or other adenine nucleotide structures and products of purine group.^32^ However, the SERS spectra of Gram-negative UTI pathogens from the family Enterobacteriacae such as *E. cloacae, E. coli and K. pneumoniae* exhibit similar spectral features with slightly shifted Raman peaks indicates that SERS-chip can be used for phylogenetic clustering of diverse bacteria. The intense Raman peak of protein at 1467 cm^-1^ for *E. coli*, nucleic acids at 1321 cm^-1^ for Klebsiella, and amide III protein at 1592 cm^-1^ for Enterobacter offers distinct features for the classification in species level. The Gram-positive *S. aureus* exhibits specific additional enhanced Raman peaks at 1005 cm^-1^ and 1180 cm^-1^ attributed to amino acids and lipids, respectively. Further, Gram-negative *P. aeruginosa* exhibits distinct peaks at 937 cm^-1^ and 1319 cm^-1^ assigned for polysaccharides and nucleic acids of the bacteria cell wall.

**Figure 4.**
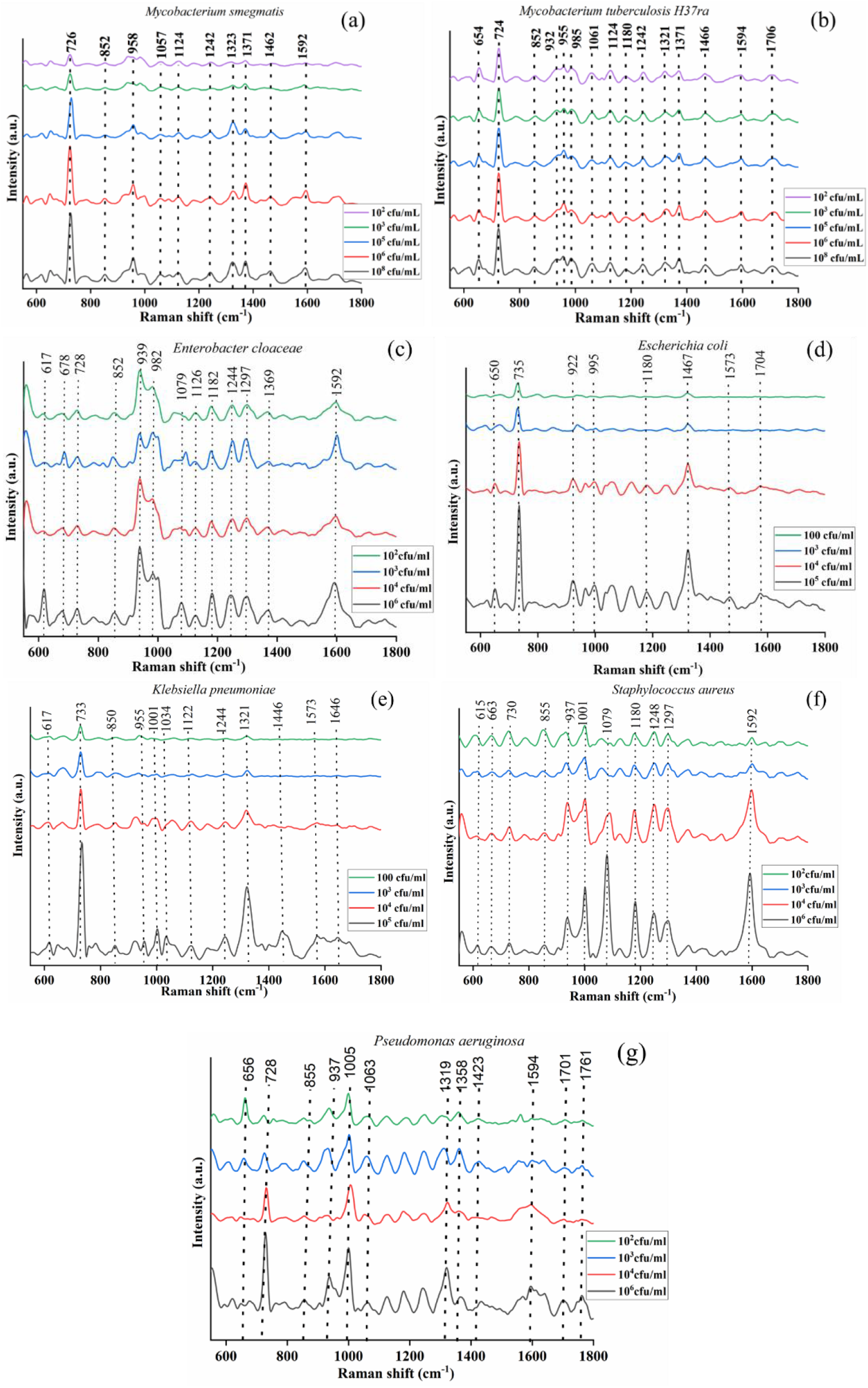
(a-g) Investigation of SERS spectra of clinically relevant bacteria from concentrations of 10^6^ −100 CFU/ml. (a, b) Tuberculosis causing bacteria and (c-f) 5 UTI-causing pathogens. The spectral signatures are unique and distinct for various species, even at lower concentrations.

Figure 3(c) shows the SERS spectral features of 3 spore forming Gram-positive plant pathogenic bacteria. The Gram-positive *Bacillus clausii* has active and reproductive spore structures whereas *Bacillus subtilis* forms endospores that have non-reproductive structures. The SERS peak at 1001 cm^-1^ is a signature of spore forming bacteria attributing due to biomarker like calcium dipicolinate from bacterial spores.^33^ The *Bacillus subtilis* has intense peaks at 939 cm^-1^ and 1601 cm^-1^ attributed to C-N stretching, and amide I protein from bacteria cell wall. The SERS peak of *Paenibacillus borealis* belonging to bacilli exhibits specific Raman peaks at 1250 and 1297 cm^-1^ attributed to nucleic acids and protein.

Figure 3(d) shows the SERS spectral features of two Gram-negative plant pathogenic bacteria *Xanthobacter autotrophicus* and *Pseudomonas citronellolis*. The SERS spectra exhibit strong SERS peak at 1003 cm^-1^ attributed to the C-C stretching of protein, as shown in Figure 3(d). This indicates the cell wall of Gram-negative bacteria is rich in protein. The reproducibility of SERS spectra for all the bacteria shown in Figure 3 was systematically performed and is presented in supplementary figure S.F. 6 in the Supplementary Section 8. The SERS characteristics of bacteria are mostly related to the vibrational modes of cell wall components such as lipids, nucleic acids, membrane proteins/lipopolysaccharides, and peptidoglycan and the metabolites of purine breakdown such as adenine, xanthine, hypoxanthine, uric acid, and guanine.^32^

Post-reproducibility investigation of SERS spectral study to determine the utility of SERS chip for identifying the TB-causing and major UTI-causing pathogens were also recorded for the dilution range of 10^6^ CFU/ml to 100 CFU/ml, as shown in Figure 4(a-g). SERS peaks in Fig. 4 show distinct and unique peak pattern for each bacterium for the entire dilution range. Even at the lowest dilution i.e. 100 CFU/ml, each bacterium retains its characteristic spectral pattern, indicating high sensitivity of Ag-SiNWs for species level detection of Raman spectra. The corresponding calibration curves of CFU/ml vs intensity count are shown in Supplementary figure S.F. 7. The calibration curve varies linearly for 4 UTI species, such as, *E. coli, K. pneumoniae, P. aeruginosa*, and *S. aureus*, and varies nonlinearly for *E. cloaceae*.

### SERS chip enables detection of Raman spectra of *E coli* in synthetic urine

To mimic urine from UTI patients, we investigated the SERS spectra of *E. coli* spiked with sigmatrix urine diluent within the range of 10^5^ CFU/ml to 100 CFU/mL, shown in Figure 5(a). The SERS peaks of urine diluent shows a distinct peak at 1001 cm^-1^, attributed to urea.^30^ The bacteria spiked urine sample exhibits two distinct enhanced peaks at 730 cm^-1^ and 1323 cm^-1^ indicating bacterial nucleic acids. The spectra collected from each dilution (10^5^ cfu /ml-100 cfu /ml) matched well with the spectra collected from the culture media.

**Figure 5.**
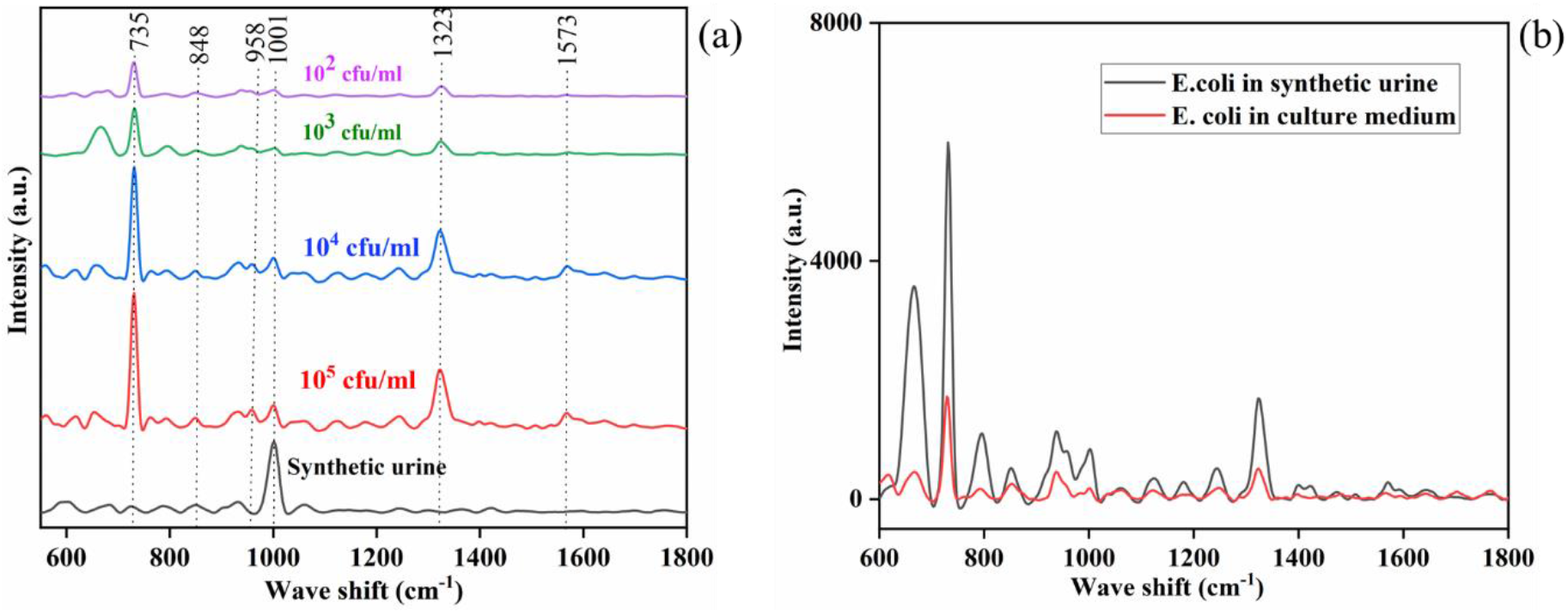
Investigation of SERS spectra of E. coli in synthetic urine media. (a) SERS spectrum of synthetic urine and E. coli bacteria at various concentrations. (b) Comparison of SERS spectra of E. coli in culture media and synthetic urine, 10^3^ CFU/ml. The SERS peak enhancement is higher when the E. coli is inoculated in urine sample.

Interestingly, we observed a significantly enhanced photon counts of SERS peaks of *E. coli* in synthetic urine as compared to the media broth, as shown in the Figure 5(b). The reason behind the substantial enhancement may have happened because of the media change influencing the bacterial contact with plasmonic nanoparticles.^30^ When *E. coli* is inoculated in a nitrogen-rich urine diluent and allowed to absorb it, the sample’s ability to bind to the SERS substrate increases significantly^30^. Hence, we can postulate that the SERS chips can detect bacteriuria.

### SERS chips enables detection of Raman spectra of resistive and susceptible strains of *E. coli*

Three different strains of *E. coli* were investigated to determine the utility of SERS-chip to capture the enhanced Raman spectra of antibiotic resistant bacteria. The *E. coli* isolate *CCUG 17620* did not possess any antibiotic resistance genes. To generate the ground truth on AMR resistance, whole genome sequencing (WGS) was obtained from previously published results and shown in Table 3.^26,27^ A wide variety of ARGs conferring resistance towards Beta-lactamase, sulfonamides, tetracyclines, and aminoglycosides was identified in both *E.coli NCTC 13441* and *A2-39*. Figure 6 illustrates the SERS spectra of various *E. coli* strains in relation to their antibiotic resistance profile. *E. coli CCUG 17620* is a wild type and *E. coli A2-39* and *E-coli NCTC 13441* are resistant to antibiotics, and the antibiotic resistance genes detected from WGS in Table 3. The WGS data was correlated with the corresponding SERS spectra. The common intense SERS peaks at 1333 cm^-1^ and 728-734 cm^-1^ correspond to the vibrational modes of adenine, polyadenine, or adenosine monophosphate of bacterial DNA. The wild-type strain of *E. coli CCUG 17620* shows a Raman peak at 1523 cm^-1^ coming from nucleic acids.^33–36^ The distinct peaks of *E. coli A239* at 1242 and 1472 cm^-1^ indicate a mixture of aromatic amino acids and nucleic acid. The SERS spectrum of AMR-resistant *E. coli* samples exhibit some additional Raman peaks at 1101 cm^-1^, 1183 cm^-1^, 1242 cm^-1^, 1472 cm^-1^, and 1623 cm^-1^. The SERS peak at 739 cm^-1^ is shifted slightly in the case of cefotaxime-resistant genes. The *E. coli NCTC 13441* exhibits several additional peaks at 1101 cm^-1^, 1183 cm^-1^, 1447 cm^-1^, and 1623 cm^-1^. The Amide III protein peak at 1280 cm^-1^ is shifted to 1242 cm^-1^ for *E. coli (A239).^34^* The low-intensity CH2 deformation peak originates from lipids at 1412 cm^-1^ of wild-type *E. coli* shifts to 1447 cm^-1^ and 1472 cm^-1^ for the resistive genes.^35^ The strain-specific features result in additional peaks that might occur due features associated with antibiotic resistance that may be unique features of a given strain. The SERS spectra exhibit the Raman peaks originating from the bacterial cell wall and components such as proteins, nucleic acids, carbohydrates, lipids, polysaccharides, and molecular complexes originating from the bacterial cell walls. The overall spectrum reveals that the strain with the greatest number of resistances exhibits extra Raman peaks. Thus, the SERS chip exhibits a distinct, enhanced, and prominent spectrum for distinct strains of a given species.

**Table 3.** Overview of bacterial strains’ phenotype and genomic background. Phenotype and genotype information from antibiotic susceptibility testing (AST) and whole genome sequencing, respectively.

| Isolate | Phenotype based on AST (antibiotic) | Plasmid presence | Antibiotic resistance genes |
| --- | --- | --- | --- |
| *E. coli* NCTC 13441 | Resistant (cefotaxime, ceftazidime, Fluoroquinolone) | Yes | blaTEM-1B, blaCTX-M-(15, 182), sul1, aadA5, catB3, dfrA17, tet(A), mph(A), qacE, sitABCD, aac(6')-Ib-cr |
| *E. coli* CCUG 17620 | Susceptive – wild type | Yes | No genes were found. |
| *E. coli* A2-39 | Resistant (cefotaxime, ceftazidime) | Yes | blaTEM-1B, blaCTX-M-2, sul1, aadA1, dfrA1, tet(A), aac(3)-Via, mdf(A), qacE |

**Figure 6.**
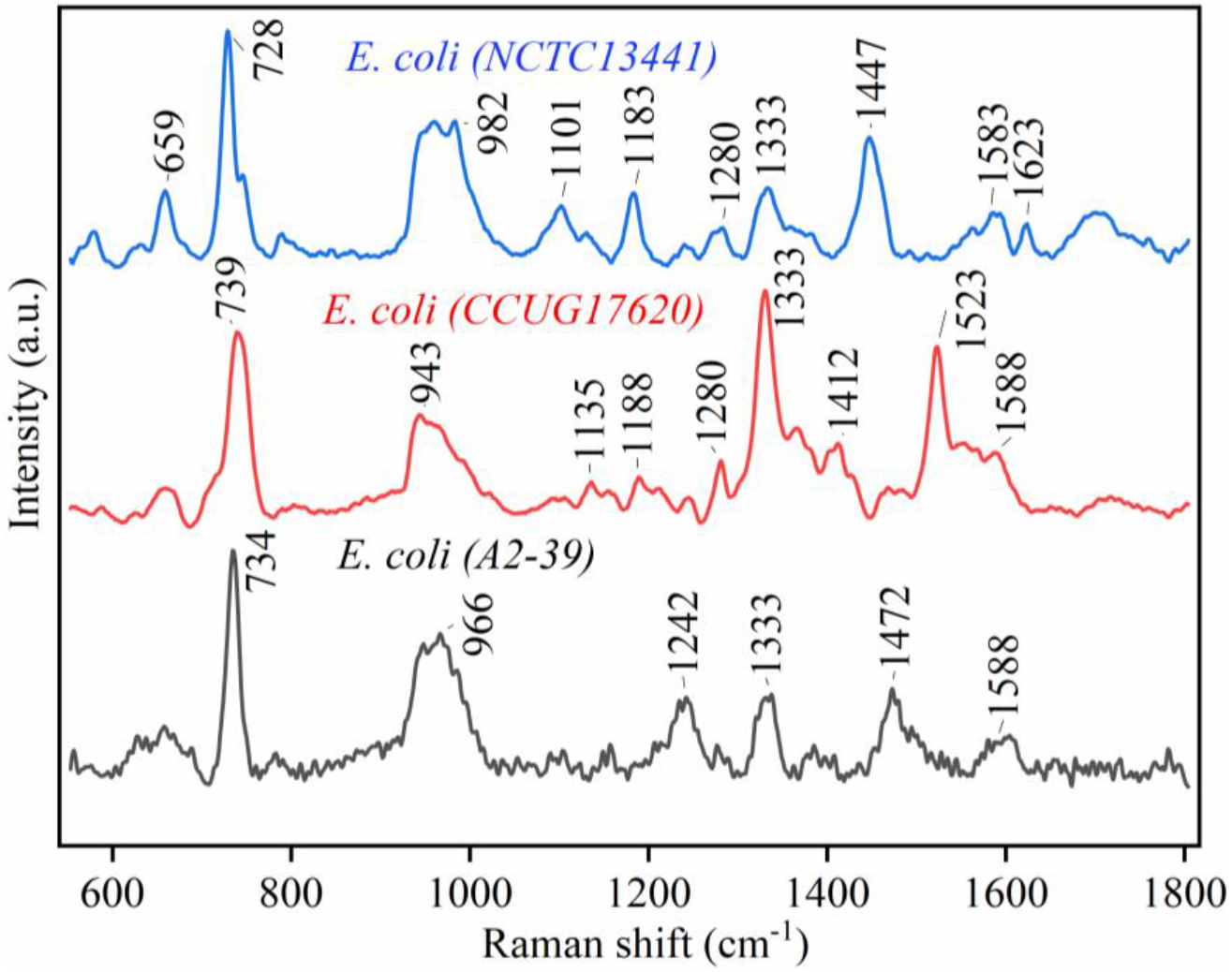
Identification of strain level of E. coli considering three different strains. Each SERS peak contains strain specific spectral information.

### Deep learning for prediction and classification of bacteria based on SERS data

The SERS data from different bacteria species using a developed model were next utilised to classification based on spectral signatures irrespective of the visual similarity among them. In total 193 separate SERS spectra were obtained from two TB causing and five UTI causing bacteria. The classification was independent of the level of dilution for a particular bacterial class. All spectra were baseline corrected using a commercially available WiRE software to reduce noise coming from the autofluorescence background. The test set was constructed using samples with varying concentrations (10^2^-10^6^ CFU/ml) extracted from each class of bacteria and the remainder was used for training. Since the model compares pairs of data for similarity during training, a new dataset was generated by pairing each training data sample with a sample from the base set. This data was then fed into the model during training.

Figure 7(a) depicts the confusion matrix for the second test set (average-performing model). We further apply the t-SNE (t-distributed Stochastic Neighbor Embedding) to the raw data and the data extracted from the flatten layer of the neural network which demonstrates that the model is able to identify features that are useful for classifying the Raman spectra of the bacteria samples. The results from t-SNE are shown in Figure 7 (b-e). Figure 7 (c) shows a tight localization for the members of the individual groups of bacteria. The performance of the trained classifier and the related confusion matrix indicate 100% accuracy in each class except for the class *M. tuberculosis* in which the classifier misclassified it as *M. smegmatis*, possibly due to the similarity between the 2 classes.

**Figure 7.**
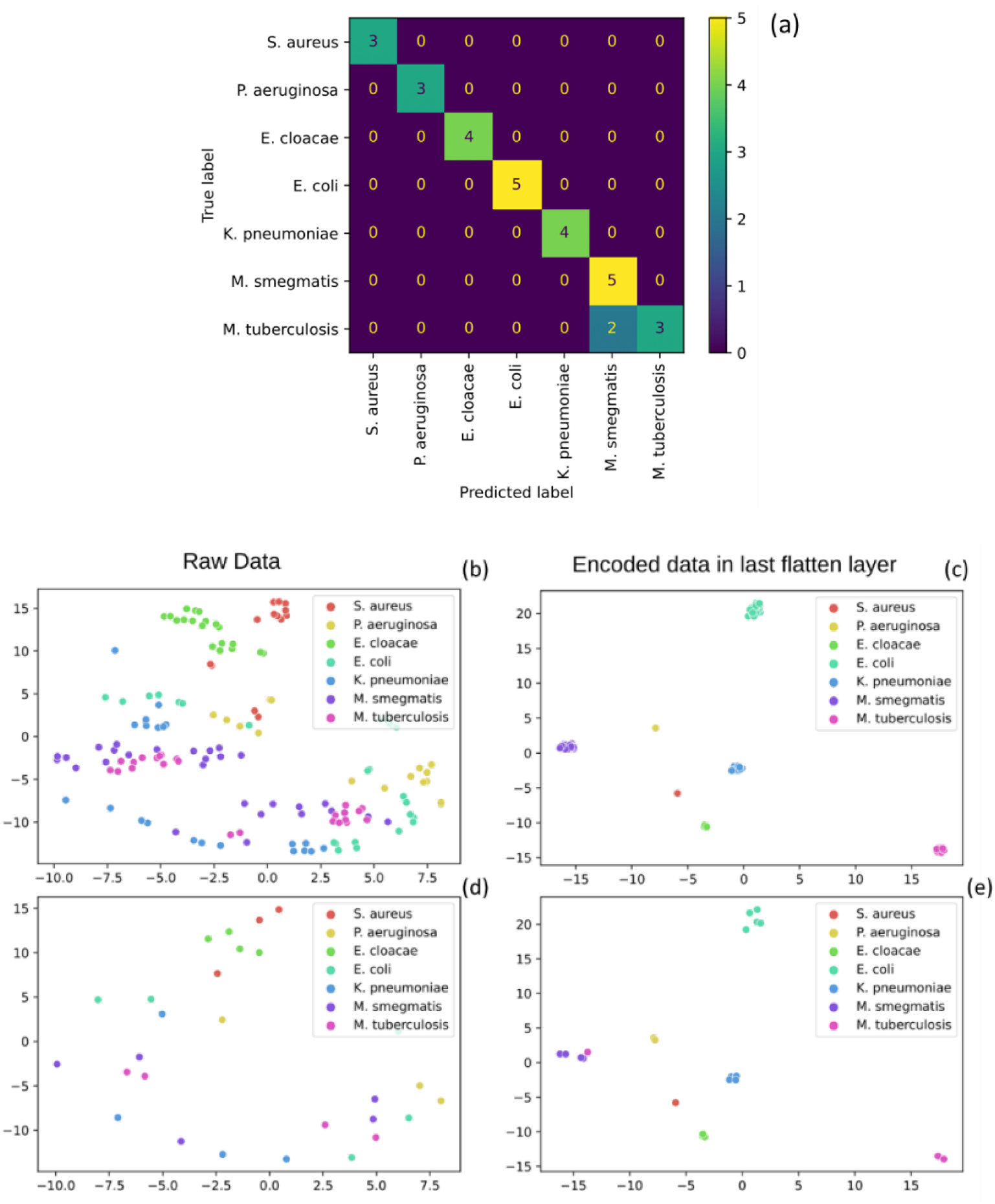
(a) Confusion matrix of the results of the second test set (average-performing model). (b-e) 2 dimensional t-SNE of raw and encoded data in the last flatten layer of the neural network, where (b,c) are training data, and (c, d) are testing data, demonstrating that the model has been able to learn distinct features and so successfully classify the bacteria from the Raman spectra.

### Classification of AMR strains using SERS data

We developed the AMR classification model using 43 samples from three distinct strains of *E.coli*, named, *E. coli* isolate *CCUG 17620, E.coli NCTC 13441* and *A2-39*. The model architecture used for the bacterial classification was also applied here. Transfer learning was used in this case by initializing the model weights with pretrained weights from the bacterial classification model.The trained model was able to obtain 100% accuracy in all three classes. By examining the t-SNE application as before Figure 7(a), we can see that the model is capable of extracting features used for the classification of the three strains, shown in Figure 8.

**Figure 8.**
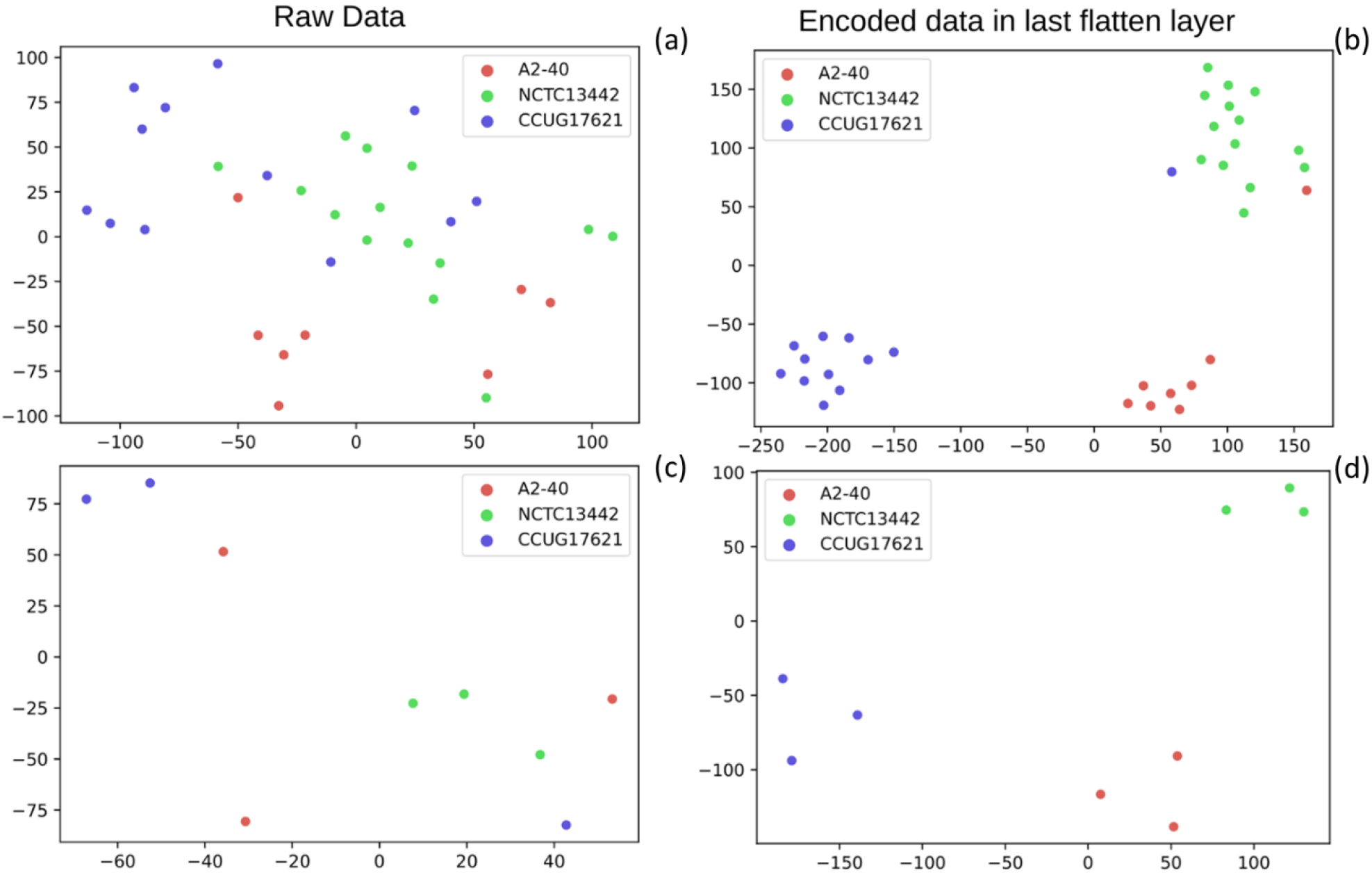
Two-dimensional t-SNE of raw and encoded data of AMR strains in the last flatten layer of the neural network, where (a), (b) are training data, and (c), (d) are testing data, demonstrating that the model has been able to learn distinct features to classify the strains of E. coli bacteria from the Raman spectra.

## Conclusions

We developed a reproducible Ag-coated Si NW SERS chip that exhibits enhanced SERS spectra of different species of bacteria with a sensitivity as low as 100 CFU/mL. The Raman spectra at the clinical threshold level of infection i.e. 10^5^ CFU/ml for UTI can be instantly obtained with excellent sensitivity using the SERS chip and portable Raman spectrometer. We demonstrated that the SERS spectra are reproducible for 12 different bacterial species and the spot-to-spot variation appears to be less than 20%. The scalability and large area fabrication method allow chips to be commercially sustainable. The trained classifier enables 100% accuracy for each class except M. tuberculosis which had two outliers that was misclassified as M. smegmatis.

The SERS data from three different strains of the same species showed strain-specific features due to molecular complexity associated with the bacterial antibiotic resistance profile. These distinct peaks are suitable for identifying strain levels information that has potential to reduce the diagnostic time and allow patients to be treated with the appropriate class of antibiotics. Thus, the Ag-coated Si SERS chips and ML can offer immediate detection of bacteria having different species and strain levels for all clinically relevant pathogens.

The utility of SERS chip to identify Raman spectral signatures of E. coli in a synthetic urine within 20 minutes after receiving the sample was demonstrated. The entire study enables identification and classification of a wide range of bacteria class in a short time. The proposed assay can be used directly in urine specimens to identify bacteria at species level along with the CFU/mL. Hands-on-time is about 5 minutes for 10 samples (i.e. 30 sec /sample). This can help the clinician to decide on whether or not to treat the patient with an antibiotic. The clinician will have information on both the infecting species and the load based on the data exploration from the CFU/mL experiments (Supplementary figure S.F. 7). In addition, this method can also be used in a microbiology lab to bypass classical biochemicals (i.e, MMTPs) used for species level identification which typically takes 24 hours after visualizing the colonies. Culture-based Kirby-Bauer Disc Diffusion or VITEK-based assays are presently used to identify pathogens associated with a bacterial infection. Ag coated Si NW SERS and the developed network model can be used for rapid label-free detection of bacteria on the site of collection or at the laboratory. In the present work a small number of samples were tested to develop the proof-of-principle. To increase the method’s reliability and for potential future use in clinical microbiology setting, the future focus will be on testing additional bacterial strains, AMR profiles, and other clinically relevant pathogens. The present investigation is based on using a single chip for a single sample at a time. The MACE technology is compatible with multiplexing, where it is possible to utilize several micro-wells for testing different samples on a single chip.

## Author Contributions

Conceptualization, Fabrication of SERS substrate, Data curation, Formal Analysis, Methodology, Software, Writing – original draft: Sathi Das, Resources, Fabrication of SERS substrate, Discussions, Visualization, Writing – review & editing: Kanchan Saxena, Jean-Claude Tinguely, Bacteria sample preparation, visualization, Writing – review & editing: Arijit Pal, Data analysis, Bacteria Classification, Machine learning, Writing – review & editing: Nima Wickramasinghe, AMR Bacterial samples, WGS, review & editing: Abdolrahman Khezri, Resources, Writing – review & editing: Vishesh Dubey, Azeem Ahmed, Supervision, Resources, Validation, Visualization: Vivekananda Perumal, Rafi Ahmad, Software, Data analysis, Classification, Writing – review & editing: Dushan N.Wadduwage, Funding acquisition, Investigation, Writing – original, review & editing: B.S. Ahluwalia, Project administration: B. S. Ahluwalia and D. S. Mehta

There are no conflicts to declare.

## Acknowledgements

Research Council of Norway funded INTPART grant nanoSymBioSys (id. 309802). Norwegian Agency for International Cooperation and Quality Enhancement in Higher Education funded AMR-Educate project (project number UTF-2020/10139). S. Das wishes to express gratitude to Laxman Prasad Goswami for his unconditional support.

