## Supplementary material for "SERS nanowire chip and machine learning enabled instant identification and classification of clinically relevant wild-type and antibiotic resistant bacteria at species and strain level": Suplementary

*[1]Bio-photonics and Green Photonics Laboratory, Department of Physics, Indian Institute of Technology Delhi, Hauz-Khas, New Delhi-110016, India*
*[2]Amity Institute of Renewable and Alternative Energy, Amity University, Uttar Pradesh, Sector-125, Noida-201303, India*
*[3]Department of Physics and Technology, UiT The Arctic University of Norway, Tromsø, 9037, Norway*
*[4]Kusuma School of Biological Sciences, Indian Institute of Technology Delhi, Hauz Khas, New Delhi 110 016, India*
*[5]Department of Electronic and Telecommunication Engineering, University of Moratuwa, Moratuwa, Sri Lanka..*
*[6]Department of Biotechnology, Inland Norway University of Applied Sciences, Holsetgata 22, 2317, Hamar, Norway.*
*7 Center for Advanced Imaging, Faculty of Arts and Sciences, Harvard University, Cambridge, USA.*

### Supplementary Section 1: Fabrication of SERS chip using MACE technique

#### Chemicals

Hydrofluoric acid (HF, 30%), Silver nitrate ($AgNO_3$, 0.1 M), Nitric acid ($HNO_3$, 70%), Hydrogen peroxide ($H_2O_2$, 30%), Sulfuric acid ($H_2SO_4$, 99.99%), and sigmatrix urine diluent. All chemicals are purchased from Sigma Aldrich.

#### Experimental protocol

A P-type Si wafer, having <100> crystal orientation and a resistivity of 5 Ω-cm, is cut into square pieces of 1 $cm^2$ area. The wafer pieces are cleaned ultrasonically using acetone, isopropanol, and deionised (DI) water. The ultrasonically cleaned pieces are kept in piranha solution ($H_2SO_4$ and $H_2O_2$ mixture (3:1)) for 30 mins to remove organic residues from their surface. Then the wafer pieces are washed with DI water and placed in a buffer HF solution to remove the native oxide ($SiO_2$) layer for 3 mins. Finally, the cleaned wafers were thoroughly washed with DI water and dried with nitrogen ($N_2$) gas.

The wafers were immersed in a mixture of 22.6 M hydrofluoric acid (HF) and 12.5 mM silver nitrate ($AgNO_3$) inside a Teflon beaker for 20 seconds to deposit over the wafer surface. The nonuniformly dispersed Ag on the Si surface changes the wafer colour from black to whitish brown. The wafers were immediately washed in DI water to remove non-attached Ag ions and placed in an etchant solution of 40% HF, 30 % $H_2O_2$ and DI water in a volumetric ratio of (2:1:1). The wafers were then kept for 90 mins inside the etchant. After the etching process, the wafer colour gets changed to grey. The next step is to place the Ag-coated etched Si wafer inside a 50% nitric acid ($HNO_3$) solution for 10 minutes to remove the Ag ions. Following this step, the wafers were placed in buffer HF for oxide layer removal. Finally, the wafers were thoroughly washed with DI water and dried in $N_2$ gas. Thus, Si nanowires are formed homogeneously with a high aspect ratio. To further decorate the formed Si NW with Ag, the Si wafers were immersed in another new solution of 40 % HF and 12.5 mM $AgNO_3$ considering the different times of 40 sec, 1 min 40 sec, and 2 min 40 sec, respectively.

### Supplementary Section 2: Morphology of SERS chip

Field emission scanning electron microscopy (FESEM, FEI Quanta 200 F SEM) was performed to investigate the morphology of coated and uncoated Si nanowire. The FESEM images of the fabricated SERS chips were visualised at 30kX magnification. The surface morphology of both coated and uncoated Si nanowires were measured using Image J software.

### Supplementary Section 3: Optimisation of SERS chip

The bare Si nanowire has an average height of 2 µm, and the deposited Ag nanoparticles have a mean diameter of 50 nm. The optimisation of the Ag coating structure on bare Si NW was performed considering the SERS spectra of 1 µM R6G solution on three different SERS chips, as shown in supplementary figure S.F. 1(a). The most enhanced Raman spectrum is obtained for an immersion time of 100 secs on the Si NW compared to 40 s and 160 secs s, respectively. The FESEM images, shown in SF. 2 (a-c) are used to explain the reason of enhancement. The number of Ag nanoparticles is less when the immersion time is 40 s, supplementary figure S.F. 1 (a). The Ag nanoparticles form dendrites of micrometre size when the immersion time is longer, such as 2 min 40 s, as seen in supplementary Figure S.F. 1 (b). Supplementary Figure S.F. 1 (c) shows the densely located anisotropic metallic cap on long Si NWs developed for 1 min 40 s coating time. Thus, the closely spaced Ag nanoparticles (NPs) are ideal for excitation of localised surface plasmon resonance (LSPR). Since large nanostructures scatter light more than they absorb it, the loosely spaced Ag dendrites do not give better enhancement than the tightly spaced MNPs. The assigned vibrational Raman peak of R6G is listed in Table 1. The EDX and X-ray photoelectron spectra (XPS) are given in the supplementary figure S.F. 3.

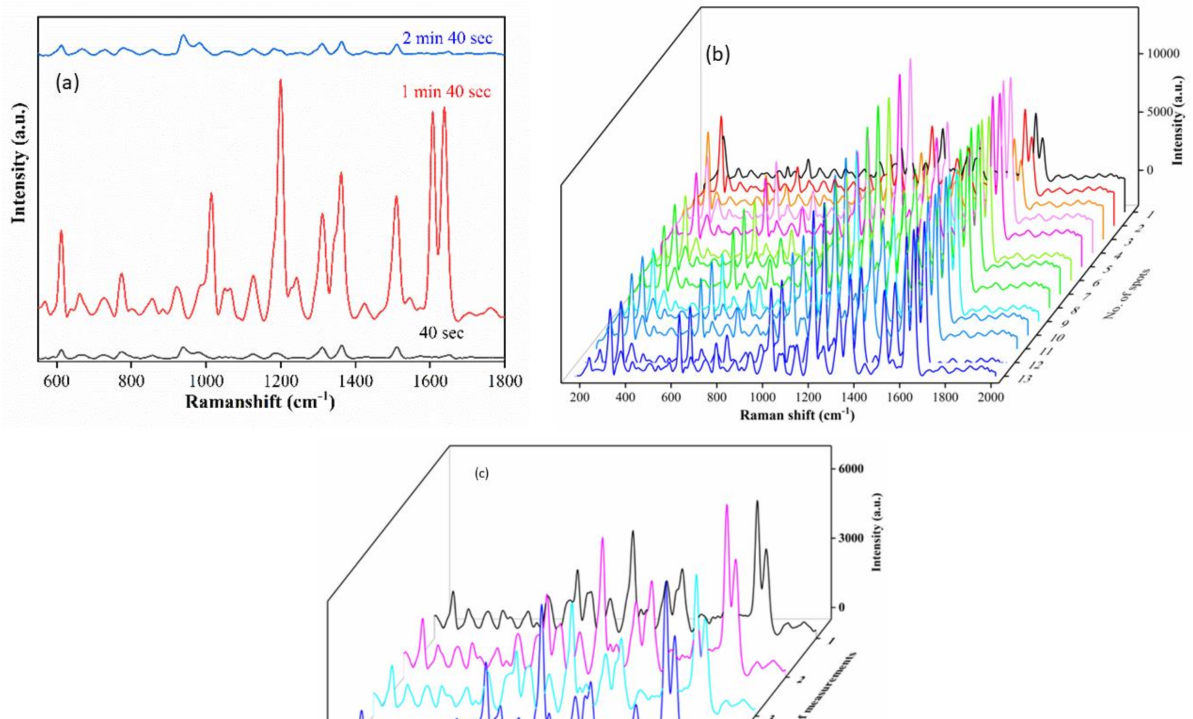


S.F. 1. (a) Optimization of the fabrication process of SERS chip. SERS spectra of 1µM R6G considering three different Ag coating times, of which1 min and 40 s provided the highest enhancement. (b, c) SERS spectra of 1 µM R6G on (b) different spots considering a particular SERS substrate, and (c) measured for four different fabrication batches, demonstrating the reproducibility of this approach.

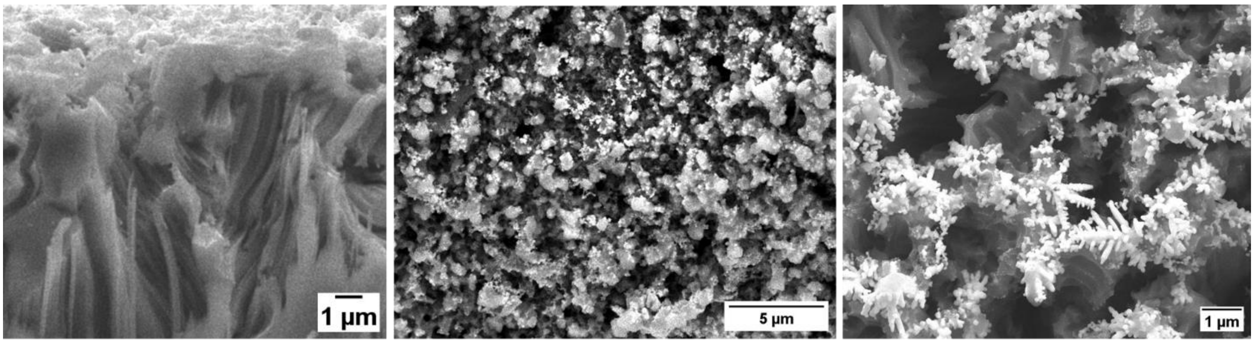


(a) (b) (c)

S.F. 2. Morphology of Ag NPs on Si nanowire for different immersion times of (a) 40 s, (b) 100 s, (c) 160 s.

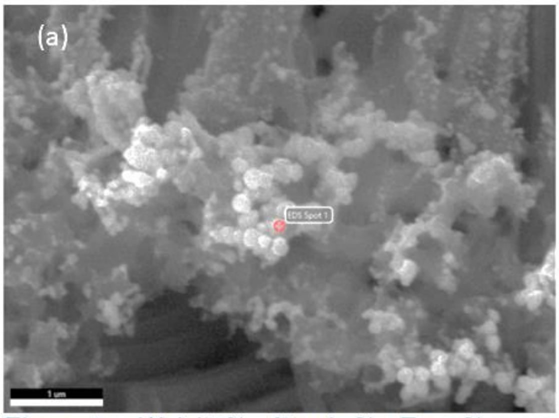


| Element | Weight % | Atomic % | Error % |
|---|---|---|---|
| Si K | 85.6 | 95.8 | 3.3 |
| Ag L | 14.4 | 4.2 | 9.2 |

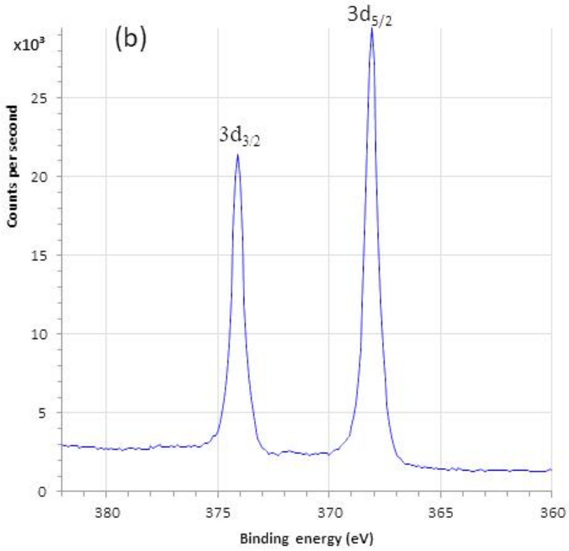


S.F. 3. (a) EDX analysis of Ag coated Si NW SERS chip. The EDX spectra confirms the presence of Ag on Si NW. (b) XPS spectra of SERS chip indicates the binding energy of Ag. The assigned peaks at $3d_{3/2}$ and $3d_{5/2}$ confirm that the deposited Ag nanoparticles are in metallic form.

**Supplementary Section 4: Finite difference time domain (FDTD) simulation of fabricated SERS chip:**

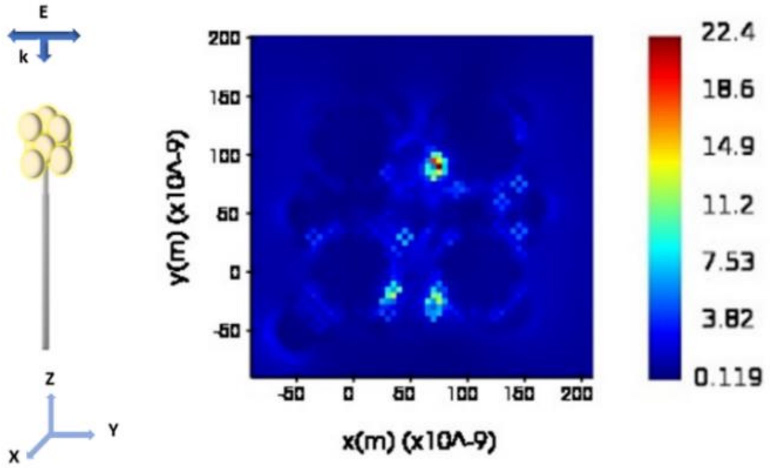


S.F. 4. (a) Geometry of FDTD simulation setup. (b) Figure shows the distribution of hotspots indicating the local electric field enhancement, the colour bar indicates $|\frac{E_{Local}}{E_{Incident}}|$.

A 3D FDTD simulation set up wass built using Lumerical (Ansys software) for the evaluation of hotspots due to nanostructures. The Si nanowire and Ag coating structures were considered for the optimised SERS parameters, extracted from FESEM images. The length and diameter of SiNW were 2 µm and 140 nm, respectively. Ag nanoparticles with a diameter of 50 nm were considered. The simulation was considered for an Ag-coated single nanowire, as shown in S.F. 4 (a), to achieve a feasible memory size. The optical properties of Si and Ag are taken from Palik data base inbuilt in the software. To reduce the memory, the mesh size was kept at 2 nm. The laser at 785 nm excitation wavelength with a band width of $\pm$ 5 nm is incident on the nanostructure plane (XY) having the k vector along the z direction. The Figure S.F. 4 (b) shows the local electric field distribution map in X-Y plane and the enhanced electric field due to the metallic Ag at the hotspot locations are evaluated to find out the amount of plasmonic enhancement.

### Supplementary Section 5: Machine learning using SERS data for bacterial classification

#### Classification strategy: Model architecture

S.F. 7. shows the model architecture used to classify the signals. The input signals initially go through four convolutional blocks. Each convolutional block is made up of a convolutional layer with an increasing number of filters and kernel size, a swish activation function, and an average pooling layer. The final convolutional layer is followed by a global average pooling layer, which produces a flattened output of size 512. The Euclidean distance between the encoded data of the base signal and the tested signal is then calculated. This value is then passed through a sigmoid function to calculate the final similarity score.

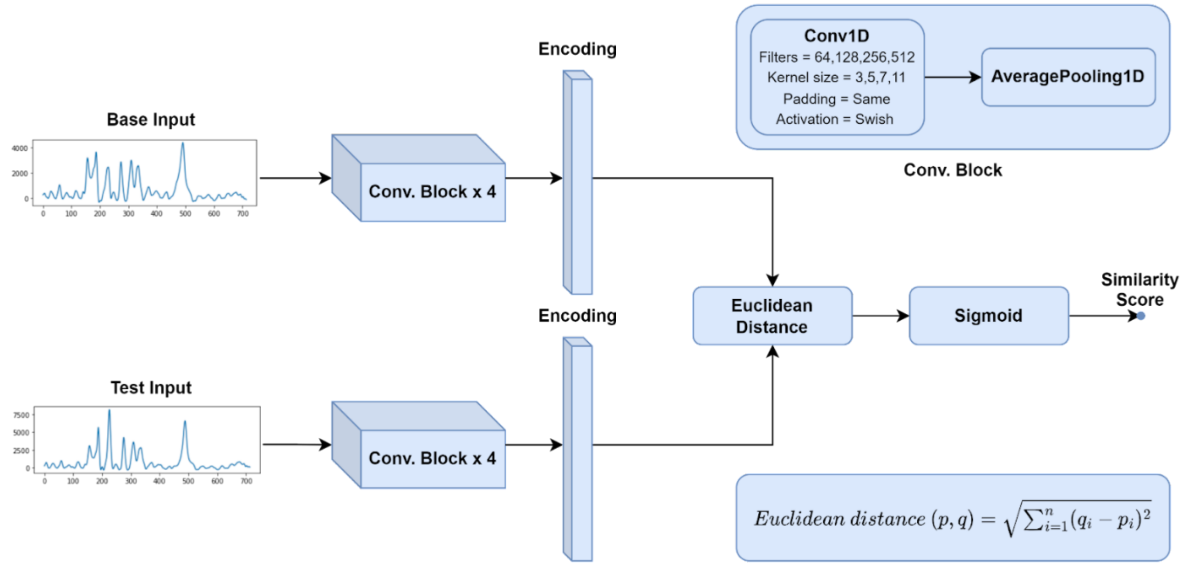


S.F. 5. Model Architecture for classification using SERS data.

#### Model inference

The signals from the test set were compared to each signal from the base set. The predicted class was determined by the signal with the highest similarity score. To check for reproducibility, three models were trained using different base, training, and test sets.

SHAP (Shapley Additive Explanations) was used for model interpretation. The SHAP values indicate how much a single feature influenced the prediction. As required by the SHAP algorithm, the training data served as the background distribution. The outputs of the third convolutional block, which are closer to the output, were used to get the SHAP values of the test data.

**Supplementary Section 6: Bacterial Sample preparation**

*Mycobacterium tuberculosis* H37Ra (ATCC 25177), *Mycobacterium smegmatis* $MC^2155$ (ATCC 700084), *Escherichia coli* K12 (ATCC 700926), *Klebsiella pneumoniae* 33495 (ATCC 33495), *Enterobacter cloacae* 10005 (ATCC 13047), *Pseudomonas aeruginosa* Migula (ATCC 27853), and *Staphylococcus aureus* 7447 (ATCC 6538P) were purchased from American Type Culture Collection (Virginia, USA). *Paenibacillus borealis* SB1 (MTCC 8085), *Bacillus clausii* 1779 (MTCC 11713), *Bacillus subtilis* 3610 (MTCC 121), *Xanthobacter autotrophicus* 10809 (MTCC 132), and *Pseudomonas citronellolis* 50332 (MTCC 1191) were procured from Microbial Type Culture Collection (Chandigarh, India). Lyophilized bacteria were revived and a loop full of revived bacteria (~10 µL) were inoculated into 1 mL of freshly prepared Luria-Bertani (LB) Broth. The primary cultures were grown overnight at 37°C with orbital shaking at 200 rpm. The primary cultures were diluted at the ratio of 1: 100 in freshly prepared Luria-Bertani (LB) Broth and grown upto mid-Log phase. The optical density at wavelength 600 nm of secondary cultures was adjusted to 0.08 (corresponds to ~ 1.0 x $10^8$ CFU/mL). Adjusted secondary cultures were serially tenfold diluted to prepare $10^6$ to 100 CFU/ml. Furthermore, during secondary culture preparation, *Escherichia coli* K12 (ATCC 700926) was spiked at the concentration of 1.0 x $10^8$ CFU/mL with sigmatrix urine diluent and then serially tenfold diluted to $10^5$ to 100 CFU/mL so as to mimic urine from UTI patients in laboratory conditions.

**Whole genome sequencing and antibiotic sensitivity profile of resistant and sensitive/wild type E. coli:**

The WGS data for *E. coli CCUG 17620*, *E. coli NCTC 13441* and *E. coli A2-39*, strains were downloaded from NCBI assembly database (BioSample: SAMN02929659, SAMEA2432036, SAMN02604091). The WGS data for isolate *E. coli A2-39* was downloaded from European nucleotide archive (PRJEB49072 and PRJEB45084). Assembly files were quality checked and further used for identifying the AMR genes and plasmid sequences as described in our previous work. [26-27]

**Supplementary Section 7: Collection of SERS spectra**

To record the SERS spectrum, a tabletop Raman spectrometer (Renishaw inVia microscope, microscope objective: 50X / 0.75 NA) and a portable Raman spectrometer Hawk Raman (New Age & Instruments) were used at an excitation wavelength of 785 nm. The spectrum acquisition power was 30 mW with 10 s integration time for all the measurements. About 5 μL R6G solution within the concentration range of $10^{-6}$ -$10^{-12}$ M were pipetted over the SERS substrate and dried in air for 24 hours before taking measurements.
A 2 μL bacterial suspensions with a serial dilution from $10^6$ CFU/ml – 100 CFU/ml were placed on the SERS chips. The SERS spectra were recorded immediately to minimize the loss of Raman signals for air-dried, photo-damaged bacteria using the portable Raman spectrometer of 785 nm excitation wavelength. However, SERS spectra for multidrug-resistant *E. coli* were generated from fixed samples (See Bacterial sample preparation section for details).

### Supplementary Section 8: Reproducibility of SERS spectra of bacteria

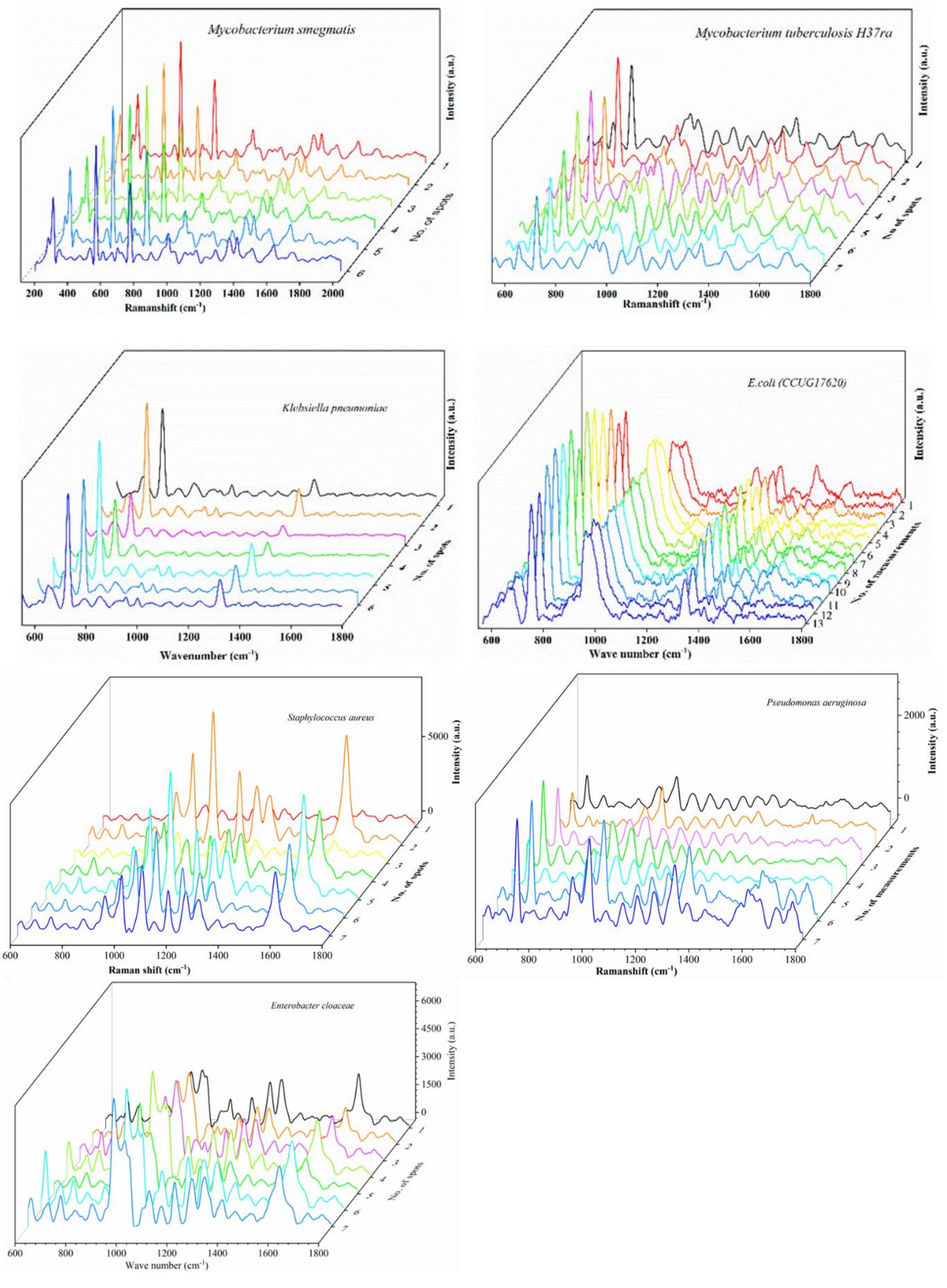


S.F. 6. SERS spectra for different bacteria at $10^5$ CFU/ml at different chip locations. The spectra visualize the reproducibility of measurements at different substrate locations.

### Supplementary Section 9: Quantification of bacteria at various concentrations using SERS spectra

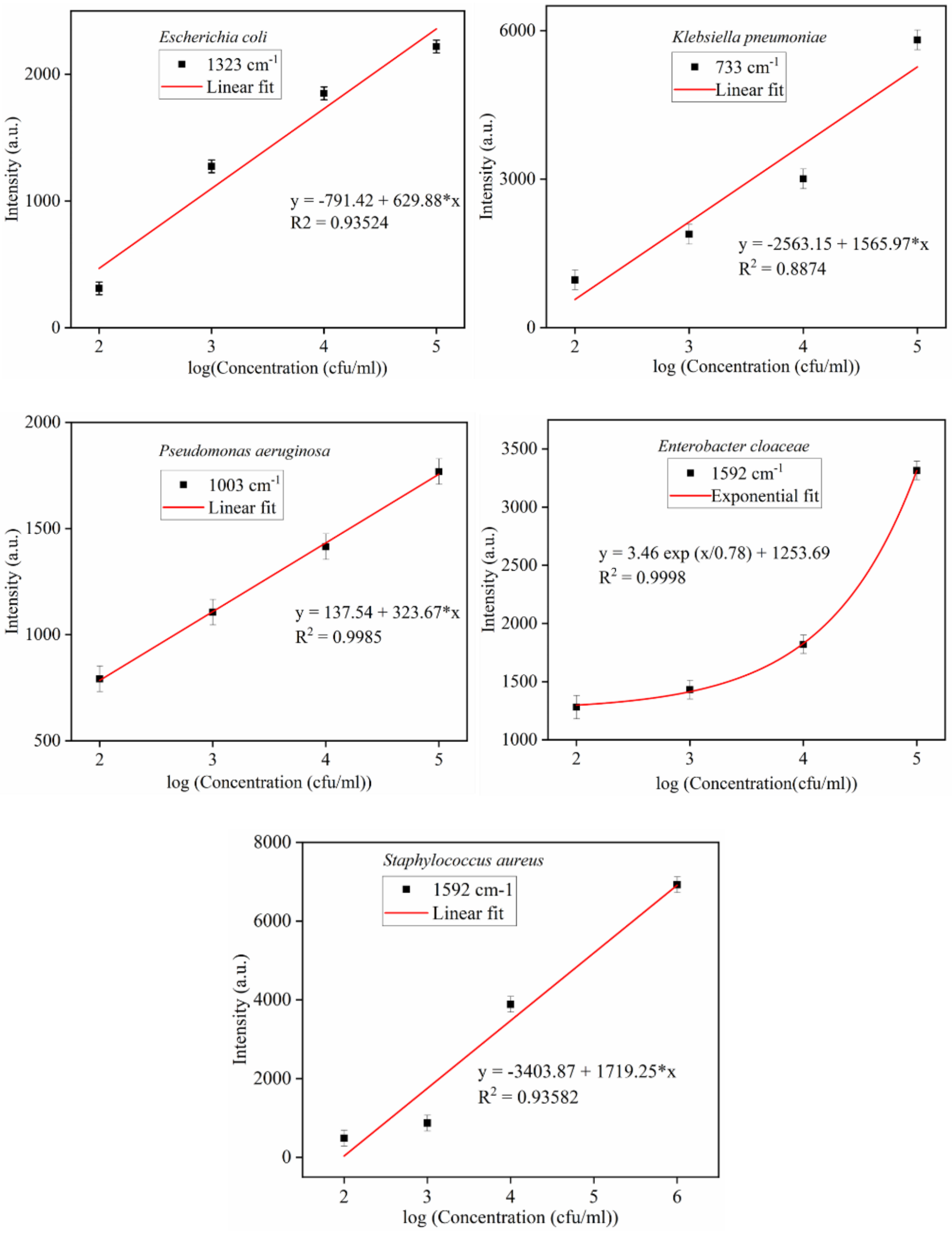


S.F. 7. Intensity vs concentration plot for five UTI causing pathogens. The calibration curve varies linearly for 4 UTI species, namely, *E. coli*, *K. pneumoniae*, *P. aeruginosa*, and *S. aureus*, and varies nonlinearly for *E. cloaceae*. Each curve depicts the variation in SERS intensity with bacteria concentration at the CFU level. The standard deviation of SERS intensity from 10 measurements over random spots for each concentration is calculated and shown in each figure. The curves could help quantify the bacteria for an unknown concentration.
